# Structure of a novel dimeric SET domain methyltransferase that regulates cell motility

**DOI:** 10.1101/264291

**Authors:** Yulia Pivovarova, Jun Liu, Johannes Lesigang, Oliver Koldyka, Rene Rauschmeier, Ke Hu, Gang Dong

**Affiliations:** Max F. Perutz Laboratories, Medical University of Vienna, Vienna Biocenter (VBC), Dr.Bohr-Gasse 9, 1030 Vienna, Austria; Department of Biology, Indiana University, Bloomington, IN 47405, USA; FH Campus Wien, Vienna, Austria

**Keywords:** AKMT, egress, lysine methylation, parasite

## Abstract

Lysine methyltransferases (KMTs) were initially associated with transcriptional control through their methylation of histones and other nuclear proteins, but have since been found to regulate many other cellular activities. The <u>a</u>pical complex lysine (<u>K</u>) <u>m</u>ethyltransferase (AKMT) of the human parasite *Toxoplasma gondii* was recently shown to play a critical role in regulating cellular motility. Here we report a 2.1-Å resolution crystal structure of the conserved and functional C-terminal portion (aa289-709) of *T. gondii* AKMT. AKMT dimerizes via a unique intermolecular interface mediated by the C-terminal TPR (tetratricopeptide repeat)-like domain together with a specific zinc-binding motif that is absent from all other KMTs. Disruption of AKMT dimerization impaired both its enzyme activity and parasite egress from infected host cells *in vivo*. Structural comparisons reveal that AKMT is related to the KMTs in the SMYD family, with, however, a number of distinct structural features in addition to the unusual dimerization interface. These features are conserved among the apicomplexan parasites and their free-living relatives, but not found in any known KMTs in animals. AKMT therefore is the founding member of a new subclass of KMT that has important implications for the evolution of the apicomplexans.

## Introduction

Lysine methyltransferases (KMTs) were first found to be involved in transcriptional control through their methylation of nuclear proteins including histones and various transcription factors [1-3]. In recent years, the roles of KMTs in other cellular activities have started to emerge [4-7]. A relatively new member of the family, the <u>a</u>pical complex lysine (<u>K</u>) <u>m</u>ethyltransferase (AKMT) was found in the phylum Apicomplexa, which includes more than six thousand species of intracellular parasites [6, 8]. Many of these parasites pose serious health threats to humans. *Plasmodium falciparum* infections claim nearly half a million lives each year [9]. Another apicomplexan parasite, *Toxoplasma gondii*, permanently infects nearly 20% of the global population. *Toxoplasma* infections in immunocompromised individuals (*e.g.* organ transplant recipients or AIDS patients) have devastating consequences, including the development of lethal toxoplasmic encephalitis due to extensive cerebral lesions [10, 11]. Similarly, maternally-transmitted *T. gondii* infection of unprotected fetuses leads to congenital toxoplasmosis [12]. To sustain an infection, the parasites need to regulate their motility appropriately in response to environmental changes to complete the lytic cycle, during which they actively invade a host cell, replicate, and disseminate by egress [11, 13-16].

Previously, we identified AKMT [for <u>a</u>pical complex lysine (<u>K</u>) <u>m</u>ethyltransferase] as a key motility regulator which controls the transition from immotile to motile behavior in the lytic cycle of *T. gondii* [6]. In immotile intracellular parasites AKMT is localized to the cytoskeletal apical complex, a set of structures at the tip of the parasite that play an important role in host cell invasion [6, 17, 18]. An influx of Ca^2+^ is the signal that normally activates parasite motility, and when cytoplasmic [Ca^2+^] increases, AKMT disperses throughout the parasite in response [6]. The AKMT knockout (Δ*akmt*) parasites generally fail to switch from the immotile to the motile state in response to this stimulus. This results in a dramatic delay in parasite dispersion during egress and a 10-fold decrease in (re)invasion efficiency, severely impairing the lytic cycle. The enzymatic activity of AKMT is important for its function in the parasite, as an enzymatically dead AKMT allele (H447V) does not complement the motility defect of the Δ*akmt* parasites. Although the native substrate(s) of AKMT are not yet known, cortical force measurement by laser trap analyses suggests that it might directly or indirectly regulate actin polymerization [19].

AKMT contains a SET-domain, which is an evolutionarily conserved structural unit that was originally identified in <u>S</u>uppressor of variegation [Su(var)3-9], <u>E</u>nhancer of zeste [E(z)] and <u>T</u>rithorax [20, 21]. Our previous phylogenetic analysis showed that AKMT orthologs form a clade close to, but distinct from, the SMYD (SET-and myeloid-Nervy-DEAF-1 (MYND)-domain) family of KMTs [22]. Importantly, while AKMT is found in all sequenced genomes of apicomplexans, there are no mammalian AKMT orthologs. This makes it a possible target for designing new, parasite-specific drugs.

In this study, we report a 2.1-Å resolution crystal structure of the structured C-terminal region of AKMT (aa289-709) without bound cofactors (apo form). The structure reveals a unique handshake-like dimer that is absent in any other SET-domain-containing KMTs, including the SMYD proteins - the nearest homologs of AKMT. Further, compared with those in the SMYD proteins, which methlylate unstructured short peptides, the putative substrate binding groove of AKMT is unusally wide, indicating that it might be capable of methylating large globular substrates. Dimerization of AKMT is mediated by a unique zinc-binding motif together with the C-terminal TPR (tetratricopeptide repeat)-like domain, which affects both co-factor binding and protein thermo stability. In *vitro* assays demonstrated that dimerization is important for the enzymatic activity of AKMT, while *in vivo* assays showed that dimerization is also required for efficient targeting to the apical complex and efficient activation of parasite motility. Taken together, our findings reveal a novel homodimeric SET domain-containing KMT that might have important implications in the evolution of a large group of parasites as well as KMTs.

## Results

### Overview of the crystal structure of *T. gondii* AKMT

The 80-kDa AKMT contains several distinct domains. Bioinformatics analyses suggested that most of the N-terminal part of the protein (aa1-300) is intrinsically disordered, while the C-terminal folded portion (aa301-709) contains four structural domains (**Fig. 1a; Supplementary Fig. 1a**). Sequence alignment indicated that the C-terminal region is well conserved among apicomplexan species, whereas the N-terminal regions is highly variable (**Supplementary Fig. 1b**). Limited proteolysis using various proteases (chymotrypsin, trypsin, thermolysin, etc.) on purified recombinant AKMT consistently generated a ∼50-kDa protease-resistant core of only the C-terminal region (data not shown). This region thus appeared to constitute the structural and functional core of the protein and was made the focus of our structural studies.

**Fig. 1.**
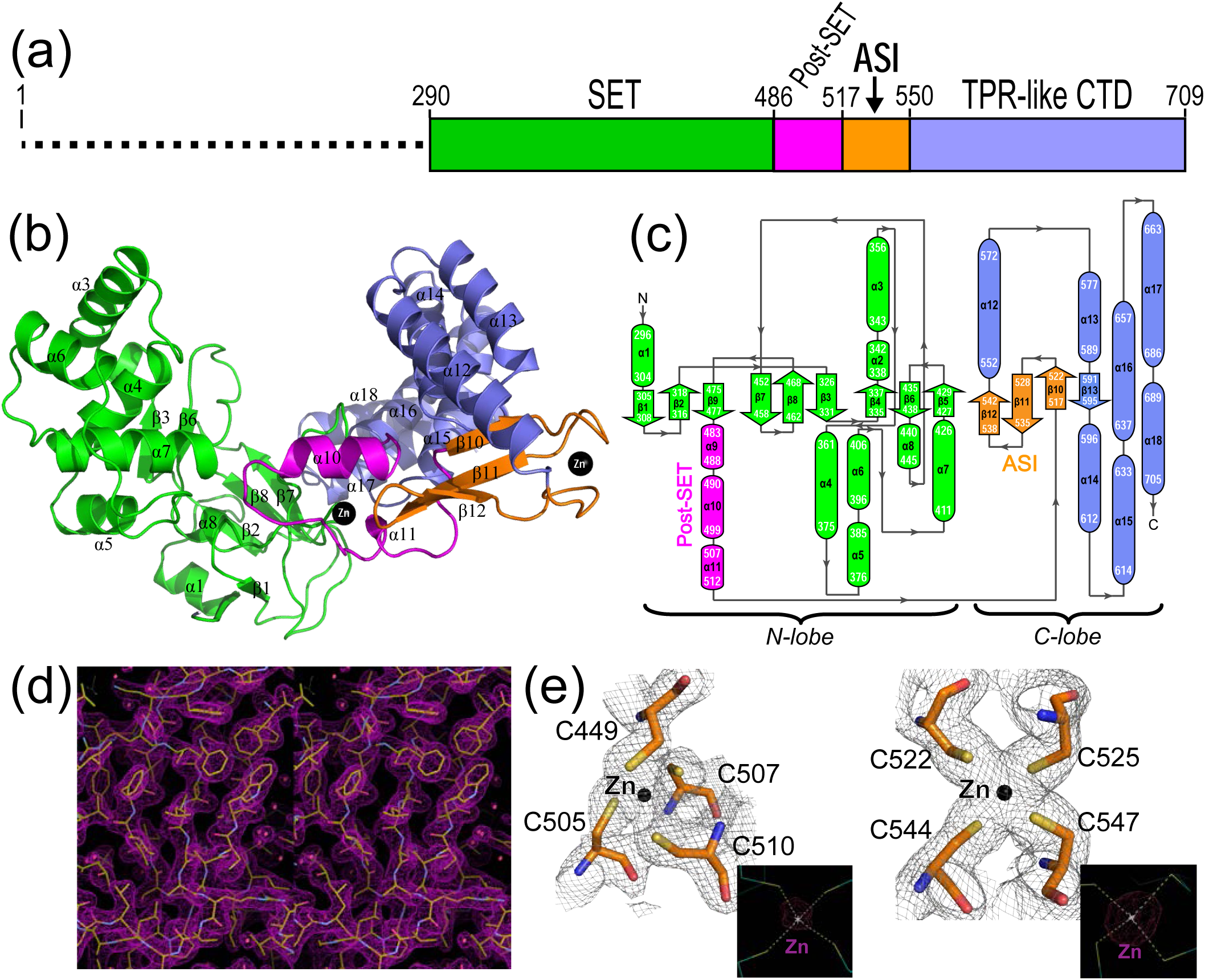
Crystal structure of *T. gondii* AKMT. (**a**) Schematic showing the domain arrangement of AKMT. The folded part used for structure determination (aa290-709) contains four predicted domains. The AKMT-specific insertion is denoted as “ASI”. (**b**) Structure of AKMT with the same color scheme as in (a). All α-helices and β-strands are labeled, and zinc ions are shown as black spheres. (**c**) Secondary structure diagram of AKMT with the same color schemes for the domains as in (a). (**d**) Stereo view of a part of the 2Fo-Fc map contoured at 1.5 σ. (**e**) 2Fo-Fc maps (1.0 σ) around the two bound zinc ions in AKMT with all cysteine residues labelled. The inserts show the anomalous difference maps contoured at 6.0 σ for the corresponding zinc ions.

A panel of N-terminally truncated AKMT constructs including aa289-709, aa295-709, and aa301-709 were expressed in bacteria. Purified recombinant proteins, both with and without reductive lysine methylation, were subjected to crystallization trials. The lysine methylated AKMT-ΔNTD (aa289-709) yielded crystals that diffracted X-rays to 2.1-Å resolution with space group *P1* (*a* = 61.94 Å, *b* = 89.37 Å, *c* = 91.72 Å; α = 108.69°, β = 101.32°, γ = 103.41°). Structure was determined using the single-wavelength anomalous dispersion (SAD) method, exploiting the anomalous signal from the bound zinc ions in the protein. The final structure had a R_work_ and R_free_ of 19.1% and 22.6%, respectively (**Table 1**).

**Table 1.** Data collection and refinement statistics.

| **Data collection** |  |
| --- | --- |
| Space group | P1 |
| Wavelength (Å) | 1.282 |
| Cell dimensions |  |
| *a*, *b*, *c* (Å) | 61.94, 89.37, 91.72 |
| $\alpha$, $\beta$, $\gamma$ (°) | 108.69, 101.32, 103.41 |
| Resolution (Å) | 20-2.1 (2.23-2.10) * |
| No. unique reflections | 167,741 (14,053) |
| *R*<sub>meas</sub> | 0.138(0.967) |
| CC(1/2) | 99.9 (79.0) |
| *I* / σ*I* | 15.5 (2.2) |
| Overall completeness (%) | 82.8 (43.2) |
| Overall redundancy | 2.8 (2.7) |
| **Refinement** |  |
| Resolution (Å) | 20-2.1 |
| Number of reflections | 167,722 |
| *R*<sub>work</sub> / *R*<sub>free</sub> (%) | 19.1/22.6 |
| No. atoms |  |
| Protein | 14,182 |
| Ligand or ion | 139 |
| Water | 434 |
| R.m.s. deviations |  |
| Bond lengths (Å) | 0.004 |
| Bond angles (°) | 0.740 |
| Ramachandran plot |  |
| Favored (%) | 96.94 |
| Allowed (%) | 3.06 |
| Outlier (%) | 0 |

\*Values in parentheses are for highest-resolution shell.

Consistent with a previous report [22], AKMT-ΔNTD was revealed to contain a SET domain (aa290-486) followed by a post-SET motif (aa487-517), together with a TPR-like C-terminal domain (CTD, aa551-709). Interestingly, the structure additionally contained a three-stranded antiparallel β-sheet (β10-12, aa518-550) between the post-SET motif and the TPR-like CTD (**Fig. 1b)**, which is absent in all other SET-domain-containing KMTs except for AKMT orthologs (**Supplementary Fig. 1b)** and thus named the <u>A</u>KMT-<u>s</u>pecific insertion (ASI) motif (**Fig. 1a and b**). AKMT-ΔNTD consisted of eighteen α helices and thirteen β strands (**Supplementary Fig. 2**), which were folded into two topologically distinct lobes (**Fig. 1c**). The final 2Fo-Fc map overall had an excellent quality (**Fig. 1d**), and the two bound zinc ions in the protein were well defined (**Fig. 1e**). Additionally, the methylated side chains of most lysine residues were resolvable (**Supplementary Fig. 3a and b**).

### AKMT forms a homodimer

The refined structural model contained four molecules per asymmetric unit, consisting of residues aa289-707, aa292-709, aa289-708, and aa289-709, which were designated as chains A, B, C and D, respectively (**Fig. 2a**). Chains A and D also contained vector-derived residues “GSHM” and “M”, respectively, at their N-termini. The final model additionally contained five sulfate (SO_4_^2-^) group, eight glycerol molecules, 21 ethylene glycols, and one diethylene glycol (**Supplementary Fig. 3c**). The four molecules in the asymmetric unit of the crystal were virtually the same with root-mean-square deviation (RMSD) of ∼0.2 Å for the Cα atoms, except for a small rigid-body shift of the N-terminal extensions between chains A/D and B/C, which is consistent with the pseudo-twofold axis related equivalence of the A-B and D-C dimers (**Fig. 2b**).

**Fig. 2.**
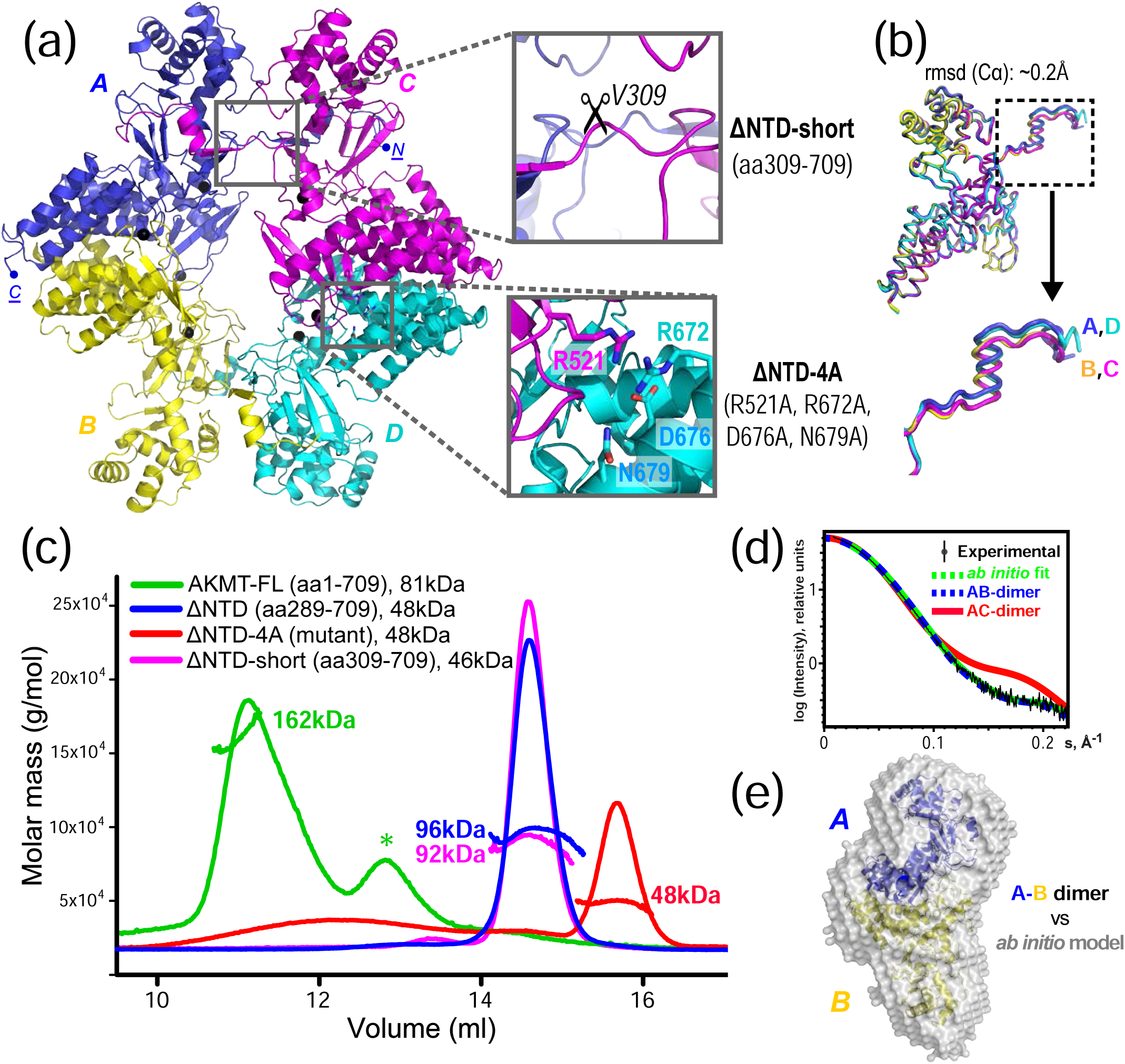
AKMT is a homodimer. (**a**) The structural model of AKMT contains four molecules (A, B, C & D, differently colored) per asymmetric unit. Enlarged boxes show the two types of intermolecular contacts. These were disrupted separately to identify the native interaction, either by deleting the N-terminal extension (ΔNTD-short, aa309-709) or by mutating the four residues that form an extensive hydrogen bond network in the vertical dimer (ΔNTD-4A). (**b**) Superposition of the four molecules in the asymmetric unit, which shows a RMSD of ∼0.2 Å for the aligned Cα atoms. The slight rigid-body shift of the N-termini (boxed) is shown in an enlarged view. (**c**) SLS data of full-length (FL), truncated, and mutated AKMT. Except for the ΔNTD-4A mutant (red), all other constructs form dimers. The asterisk (*) marks impurities co-purified with the full-length protein. (**d**) Low-angle region of the SAXS curve of AKMT-ΔNTD (black) overlaid with theoretical scattering curves for the N-terminally mediated A-C dimer (red) and the C-terminally mediated A-B dimer (blue) calculated in CRYSOL. The green curve represents the simulated scattering from *ab initio* model computed in DAMMIF. (**e**) The *ab initio* low resolution model reconstructed in DAMAVER by selecting and averaging the most probable models generated in 10 cycles of DAMMIF. The mean normalized spatial discrepancy (NSD) is 0.538. The N-terminal fragments (aa289-309) in the A-B dimer were rebuilt after rearranging the swapped domains. The model is shown superimposed on a ribbon diagram of the A-B dimer.

The tetramer in the final model had four intermolecular dimeric interfaces of two types, with the A-B dimer being structurally equivalent to the D-C dimer, and the A-C dimer equivalent to the D-B dimer. The two types of dimeric interface were both pronounced and had similar buried surface areas (BSA) of ∼1800 Å^2^ and Gibbs free energy (Δ^i^G) of approximately -25 kcal/mol (**Supplementary Fig. 4**); it was unclear whether either or both of these interfaces is physiologically relevant. To reveal the native oligomerization state of AKMT, we carried out static light scattering (SLS) analyses on both full-length AKMT and AKMT-ΔNTD. The results showed that both proteins formed homodimers (**Fig. 2c**, green and blue traces), suggesting that a dimeric conformation is the native state of AKMT. Therefore, only one of the two dimeric interfaces present in the tetrameric crystal structure is natively formed.

To determine which of the two dimeric interfaces (A-C/D-B or A-B/D-C) in the crystal structure represents the native conformation, we first disrupted the A-C/D-B dimer interface by deleting the N-terminal segment (i.e. α1 and β1) of the structure (designated as AKMT-ΔNTD-short, aa309-709) (**Fig. 2a**, upper insert). The resulting protein formed a similar dimer to the wild-type (**Fig 2c**, magenta trace), suggesting that this interface is not the native inter-dimer interface. Based on our analyses of the intermolecular interactions in the crystal structure using PDBePISA [23], eight of the ten direct hydrogen bonds and all salt bridges in the A-B/D-C interface were mediated by the side chains of three charged residues (R521, R672, and D676) plus the polar residue N679 (**Supplementary Fig. 4**). We therefore mutated all these residues to alanine to try to disrupt the interaction between chains A and B (**Fig. 2a**, lower insert). The resulting mutant (designated as AKMT-ΔNTD-4A: R521A/R672A/D676A/N679A) was found to become monomeric in SLS analyses, indicating that the A-B interface is indeed the native dimeric interface (**Fig. 2c**, red trace). For convenience, the construct AKMT-ΔNTD-4A will be referred to as the monomeric mutant in this study. To further verify the dimeric conformation of AKMT, we additionally assayed AKMT-ΔNTD in solution using small-angle X-ray scattering (SAXS). The results showed that the theoretical scattering of the A-B dimer was in excellent agreement with the experimental data (χ^2^ = 1.59), whereas the simulated curve of the A-C dimer was not (χ^2^ = 10.99) (**Fig. 2d**).

Furthermore, the A-B dimer fitted perfectly into the calculated *ab initio* model reconstructed from the SAXS data (**Fig. 2e**). Therefore, we concluded that AKMT forms a homodimer in its native state, and the dimerization is mediated by its C-terminal region. The N-terminally mediated dimerization in the crystal structure was likely an artifact caused by domain swapping during crystallization.

### The dimeric interface of AKMT contains a unique zinc-binding motif

Both the post-SET and the ASI motifs in AKMT bound a zinc ion, and the zinc ion in each case was coordinated by a cluster of four cysteine residues, C449/C505/C507/C510 in the post-SET motif and C522/C525/C544/C547 in the ASI motif (**Fig. 3a and b**). Notably, one of the cysteine residues (C449) participating in zinc binding in the post-SET motif is located in the SET domain. The ASI motif is mostly buried at the A-B dimeric interface, and mutating these zinc-binding cysteine residues in the ASI made the protein insoluble (data not shown). This suggests that the ASI motif plays a critical role in maintaining the structural stability of AKMT.

**Fig. 3.**
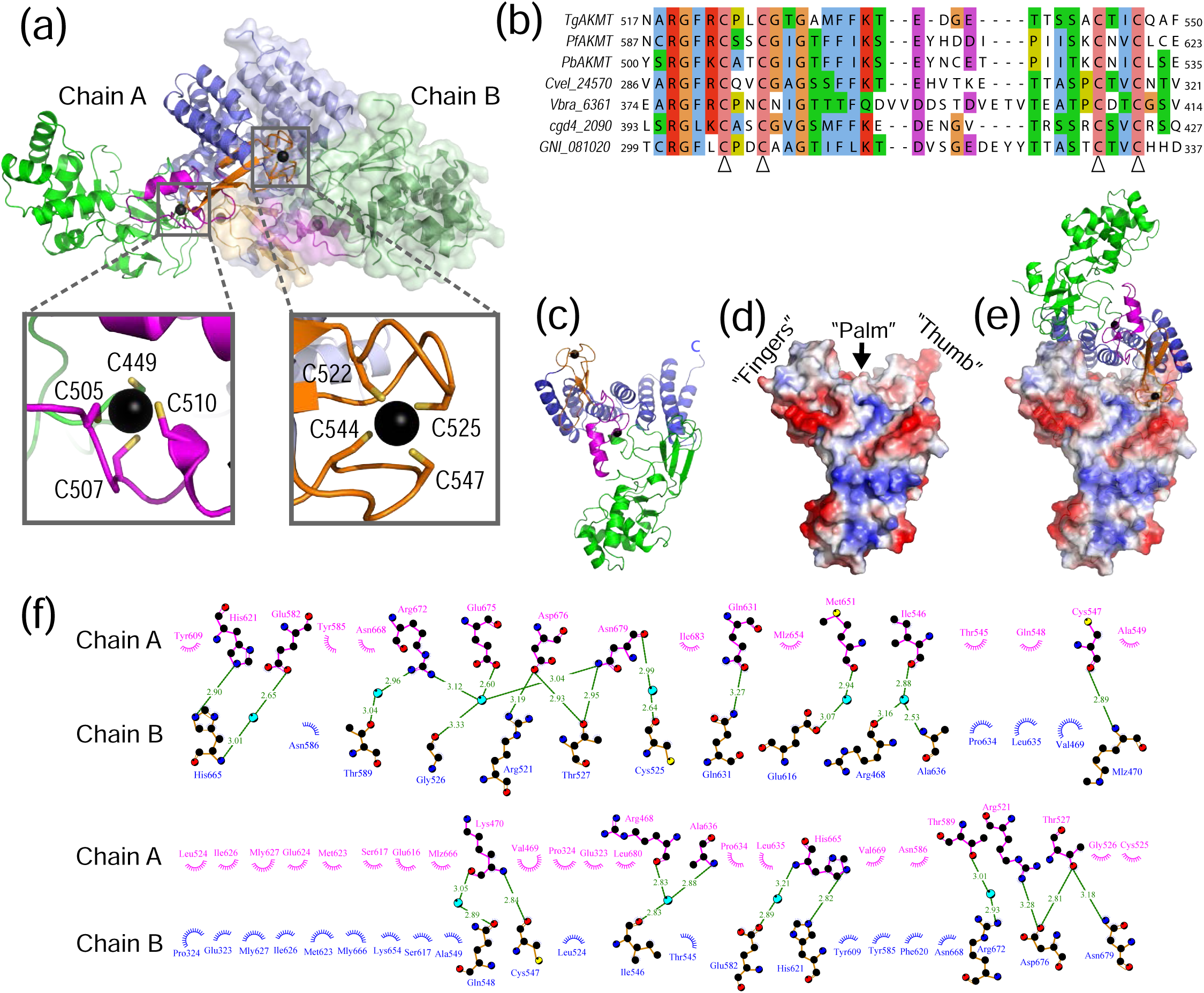
Intermolecular interactions in the AKMT homodimer. (**a**) Homodimer of AKMT with chain A shown in ribbon diagrams and chain B in a semi-transparent surface plot. Insets show the enlarged views of the two bound zinc ions in chain A. (**b**) Sequence alignment of the ASI motifs in AKMT and orthologs. The four conserved cysteines that coordinate zinc binding are indicated with open arrowheads. (**c**) Ribbon diagram of AKMT monomer with individual domains colored as in (a). (**d**) Electrostatic plot showing the hand-like structure with the C-terminal end of the protein forming the “thumb”, the ASI motif forming the “fingers”, and the central helices of the TPR-like domain forming the “palm”. (**e**) “Handshake”-like dimer of AKMT. (**f**) Intermolecular interactions in the AKMT homodimer. Residues contributing to dimer formation are plotted with DIMPLOT in the LigPlot plus suite [66]. Side chains of residues involved in hydrogen bond formation are shown as ball-and-stick. Oxygen, nitrogen and carbon atoms are colored in red, blue, and black, respectively. Water molecules mediating inter-molecular hydrogen bond formation are shown as cyan-colored spheres. Green dotted lines indicate hydrogen bonds, with the length of the bonds shown on top of the lines. Non-bonded residues involved in hydrophobic interactions are shown as spoked arcs.

A closer examination of the structure revealed that AKMT formed an “open hand”-like structure, with the ASI motif forming the “fingers”, and the central and C-terminal parts of the TPR-like domain forming the “palm” and the “thumb”, respectively (**Fig. 3c and d**). In the homodimer, the two hands were orientated in a handshake manner, with the “fingers” of one hand reaching the base of the “thumb” on the other and the two “palms” directly facing each other (**Fig. 3e**). The homodimer of AKMT was stabilized by more than twenty hydrogen bonds (directly or mediated by water molecules), two salt bridges, and many hydrophobic interactions (**Fig. 3f**; **Supplementary Fig. 4**).

### AKMT binds cofactor in submicromolar range

The lysine methyltransferase activity of AKMT has been previously tested *in vitro* using an artificial histone substrate and a ^3^H-labeled cofactor, S-adenosyl-L-methionine (SAM) [6, 22]. These assays demonstrated that AKMT is an active lysine methyltransferase capable of binding SAM, which serves as a methyl group donor in the methylation reaction. The crystal structure of AKMT reported here was obtained by crystallizing AKMT in the absence of cofactors. To verify the binding of AKMT to cofactors and to understand how the interaction occurs, we first carried out isothermal titration calorimetry (ITC) experiments. Recombinant full-length AKMT bound the cofactor SAM with a dissociation constant (K_d_) of 0.62 μM; AKMT-ΔNTD showed a similar affinity of K_d_ = 0.63 μM (**Fig. 4a**). AKMT-ΔNTD bound the cofactor analog sinefungin (SFG) with a higher affinity (K_d_ = 0.26 μM) (**Fig. 4a**), likely due to extra hydrogen bonds formed between the additional amide protons on SFG and surrounding residues of AKMT. To verify the results, multiple extra ITC measurements were performed for all above interactions as well as for the interaction between full-length AKMT and SFG, which show a consistent trend (**Supplementary Fig. 5**).

**Fig. 4.**
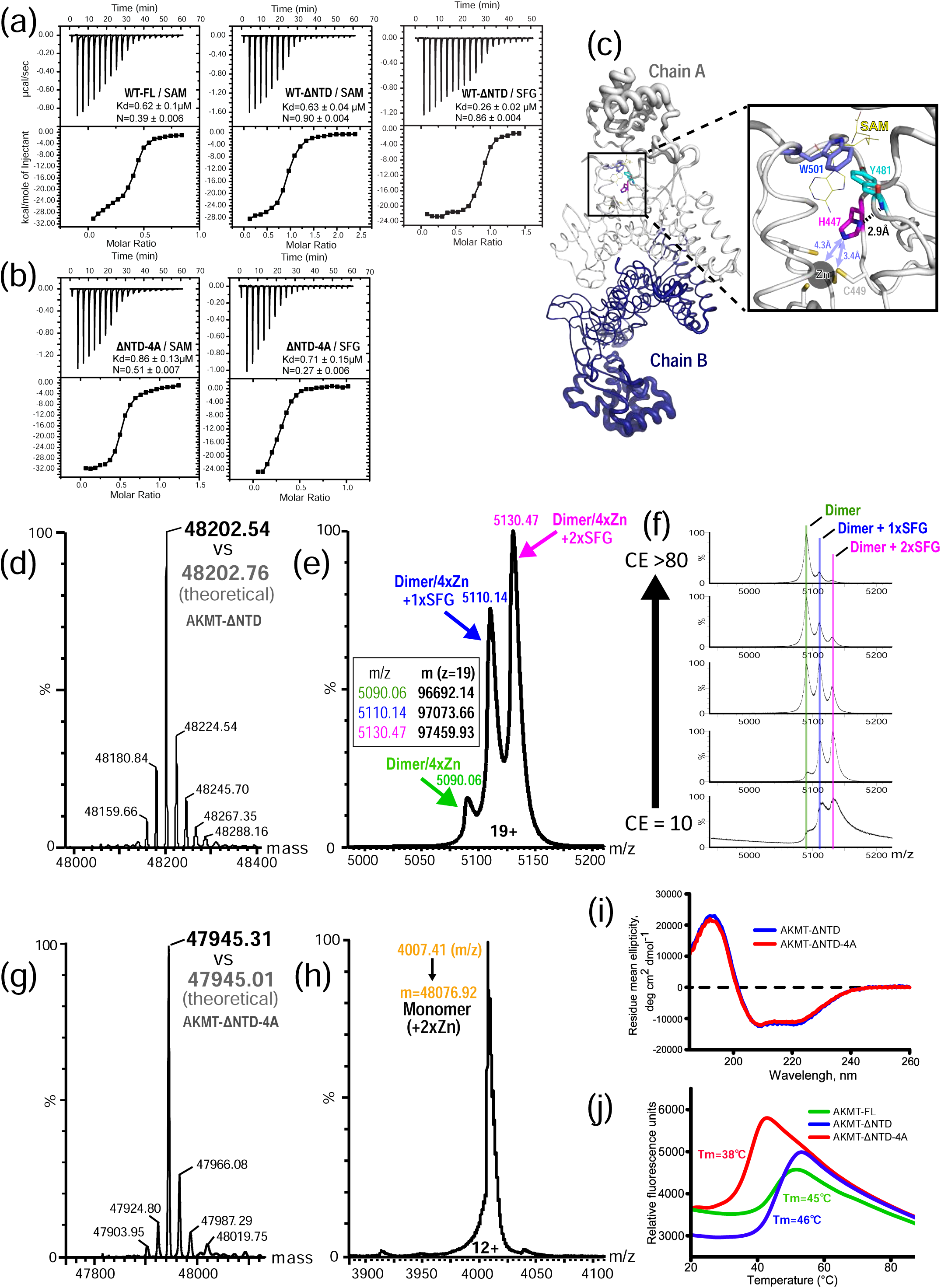
Cofactor binding and biophysical characterizations of AKMT. (**a**) ITC binding curves of wild-type full-length and AKMT-ΔNTD with the cofactor SAM or the analog SFG. (**b**) ITC binding curves of AKMT-ΔNTD-4A with SAM or SFG. (**c**) Pymol B-factor putty representation of the AKMT dimer. Width of the tubes, which is proportional to the values of the B-factors of each residue, indicates the dynamics of the local structure. Shown in the inset is an enlarged view of the highly rigid region around residue H447 that bridges the putative cofactor binding site to the zinc-bound ASI motif. The cofactor SAM (shown in yellow sticks) is based on superposition of SAM bound SMYD2 structure (5ARG.pdb) onto the AKMT structure. (**d**) Deconvoluted MS spectrum of AKMT-ΔNTD in the denatured state. The measured molecular mass (48202.54 Da) is in excellent agreement with the theoretical value calculated from the primary sequence of the protein (48202.76 Da). (**e**) Mass determination of SFG-bound AKMT-ΔNTD at the native state. Based on the calculated masses, each of the three peaks could be mapped to a subspecies of the complex, with the majority being the homodimer loaded with four zinc ions and two SFG molecules (2×SFG). (**f**) Collision-induced dissociation of native SFG-bound AKMT-ΔNTD analyzed by tandem mass spectrometry. Increasing trap collision energy (CE) led to dissociation of SFG from the dimer, but did not affect stability of the zinc-bound dimer. (**g**) MS spectrum of AKMT-ΔNTD-4A mutant acquired under denaturing conditions. Measured mass of 47945.31 Da closely matches the calculated value, 47945.01 Da. (**h**) Native MS analyses of AKMT-ΔNTD-4A in the apo form proves the monomeric state of the mutant. (**i**) CD spectra of AKMT-ΔNTD and AKMT-ΔNTD-4A. (**j**) Melting curves of various AKMT constructs by the DSF assay. AKMT-FL and -ΔNTD have a similar Tm, whereas that of the monomeric mutant is substantially lower.

The monomeric mutant AKMT-ΔNTD-4A bound SAM and SFG with K_d_ values of 0.86 μM and 0.71 μM, respectively (**Fig. 4b**). The mutant therefore showed modestly reduced cofactor binding affinity compared to AKMT-ΔNTD based on the measured K_d_ values. However, the N values for both mutant/cofactor interactions (N = 0.27 and 0.51 for interaction with SAM and SFG, respectively), which represent the molar ratio between the cofactor and the target protein, were much lower than those of the AKMT-ΔNTD (N = ∼0.9). This was probably because some molecules of the monomeric mutant in the ITC reaction chamber were unable to bind cofactors. The results were confirmed by additional independently performed ITC measurements (**Supplementary Fig. 5**).

Comparison of the structure of AKMT with its structural homolog SMYD2 in complex with the cofactor SAM revealed a putative cofactor binding site in AKMT (**Supplementary Fig. 6a and b**). In the AKMT structure we observed that the residue H447, which is highly conserved across the phylum Apicomplexa, not only forms a hydrogen bond via its side chain with the backbone amide proton of the neighboring residue Y481, which helps define the cofactor binding pocket, but is also in close proximity (3.4-4.3 Å) to the zinc-binding site in the ASI motif (**Fig. 4c**). Mutation of H447 to valine (H447V) would be predicted to disrupt its interaction with Y481, and thus perturb the structure around the putative cofactor binding site. This analysis explains our previously published observation that the H447V mutant completely abolished the enzymatic activity and *in vivo* function of AKMT [6]. Such mutation possibly also destabilizes the nearby ASI motif that forms a part of the dimerization interface of AKMT.

The stoichiometry for SFG:AKMT-ΔNTD in the ITC experiment was approximately 0.9:1, indicating that both AKMT molecules in the homodimer are capable of binding cofactors. As an additional test, we carried out mass spectrometry (MS) experiments. The molecular mass of AKMT-ΔNTD measured at a denaturing condition was in excellent agreement with the theoretical molecular weight calculated based on the primary sequence (**Fig. 4d**). Further MS analyses of SFG-bound AKMT-ΔNTD at native state showed that the majority of the protein was a dimer loaded with two SFG molecules plus four zinc ions (**Fig. 4e**). Subsequent tests for collision-induced dissociation of ligand from native SFG-bound AKMT-ΔNTD, which were analyzed by tandem mass spectrometry, revealed that increasing trap collision energy (CE) led to dissociation of SFG from the dimer, but affected neither the stability of the dimer nor its binding to zinc ions (**Fig. 4f**). Additionally we performed MS analyses on AKMT-ΔNTD-4A mutant in the apo state. MS results under denaturing conditions verified the mutant by a perfectly matched molecular mass (**Fig. 4g**), whereas native MS data confirmed the monomeric state of the mutant with two stably bound zinc ions (**Fig. 4h**).

### Dimerization of AKMT is important for maintaining its thermostability

The lower molecular ratio (*i.e.* N values) and less stable cofactor binding of the monomeric mutant compared to the AKMT-ΔNTD protein suggested that the mutant is possibly misfolded and/or structurally unstable. To test these possibilities, we first carried out circular dichroism (CD) analysis for rapid determination of the folding properties of the purified proteins [24]. The results showed similar curves for the wild-type and the monomeric mutant, suggesting that the mutant is folded similarly to the wild-type (**Fig. 4i**).

We further checked the stability of the monomeric mutant by differential scanning fluorimetry (DSF), a technique that generates a melting curve for a target protein by monitoring the increase in the fluorescence of a dye binding to accessible hydrophobic portions of the protein while the protein gradually unfolds in increasing temperature [25]. The results showed that wild-type AKMT-ΔNTD had a similar melting temperature (Tm) to the full-length AKMT (46°C vs 45°C), but the monomeric mutant had a drastically reduced Tm (38°C) compared to the wild-type protein (**Fig. 4j**). These results demonstrated that disruption of the dimerization interface renders the protein structurally less stable.

### Dimerization of AKMT is important for its function *in vitro* and *in vivo*

To test the effects of dimerization on the *in vitro* methylation activity, we compared the activities of the AKMT-ΔNTD and AKMT-ΔNTD-4A recombinant proteins in methylation reactions (**Fig. 5a**). The reactions constituted time series of four 2-fold dilutions of the enzyme from 120 nM to 15 nM. Each reaction also contained 4.1 μM of the artificial substrate, human histone H3.3, and 9.2 mM of the tritium labeled co-factor, ^3^H-SAM. 0.1mg/ml BSA was included as a carrier protein for preventing non-specific protein sticking to the test tube. The difference between the two truncations was pronounced in all enzyme concentrations tested, which shows that AKMT-ΔNTD-4A is less active than AKMT-ΔNTD as a methyltransferase.

**Fig. 5.**
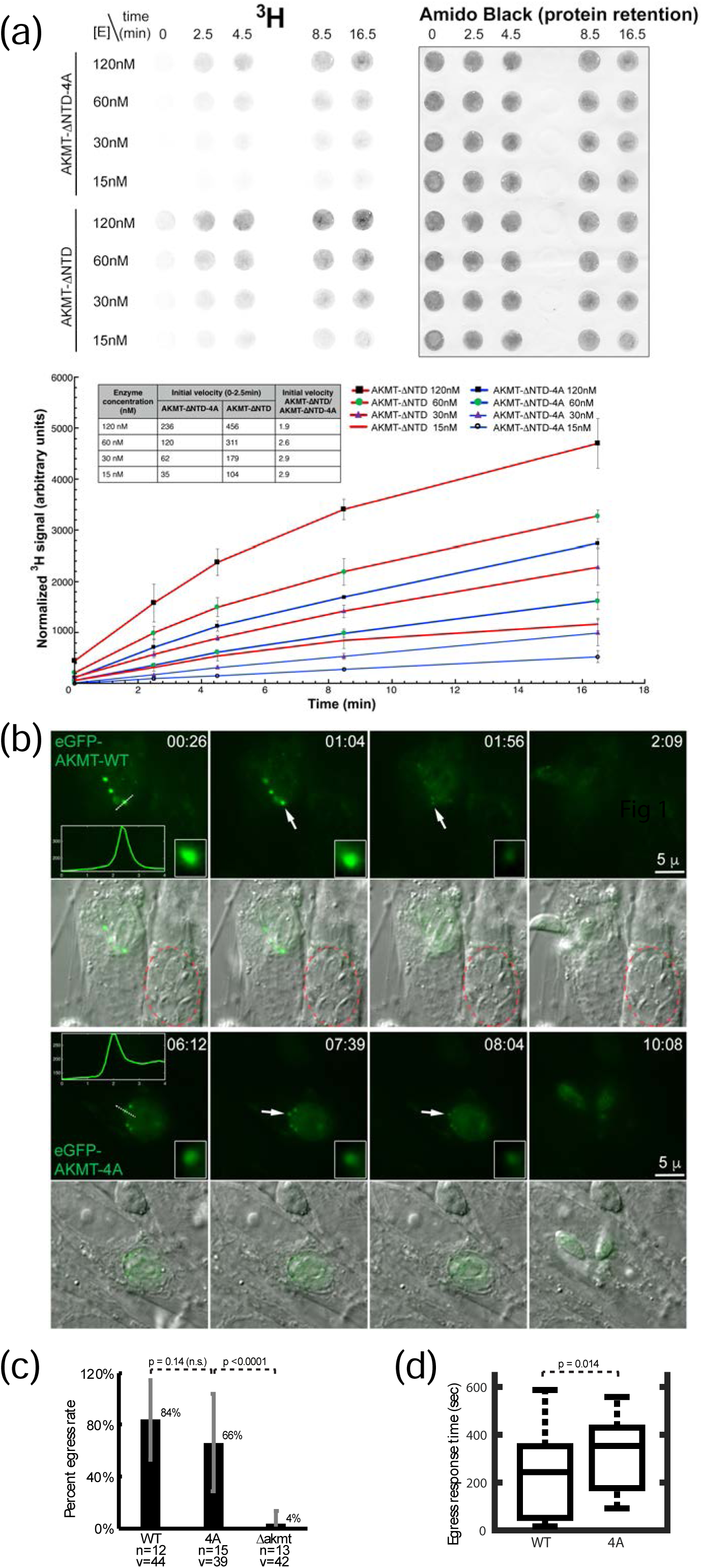
Functional defect of AKMT-ΔNTD-4A and AKMT-ΔNTD revealed by *in vitro* methylation and *in vivo* egress assays. (**a**) Methylation assay to compare the activity of AKMT-ΔNTD-4A and AKMT-ΔNTD. Top panels: Dot-blot of tritium signal (left) and total protein staining by amido black (right). Bottom panel: X-axis shows time (in minutes) of reactions spent at room temperature. Y-axis shows “normalized tritium signal” (in arbitrary units). To generate the normalized tritium signal, the signal from the Typhoon scan of the phosphorimager was first normalized against the corresponding protein staining by amido black to correct for variations in protein retention on the dot-blot to generate “retention corrected tritium signal”. Individual retention corrected tritium signals were then normalized against the sum of corrected signals of reactions in the same experiment. The error bars represent ±2 standard deviations of three independent experiments. (**b**) WT: Δ*akmt* parasites expressing eGFP-AKMT-WT. 4A: Δ*akmt* parasites expressing eGFP-AKMT-4A. Images selected from time-lapse imaging of calcium ionophore (A23187)-induced egress of Δ*akmt* parasites expressing eGFP-AKMT-WT (top panels) or eGFP-AKMT-4A (bottom panels). Inset images (4X) include the apical region from which the eGFP-AKMT-WT or eGFP-AKMT-4A translocates after the addition of A23187. Inset graphs are line plots of integrated intensity over the region indicated by the dotted lines (X-axis: distance in microns; Y-axis: intensity). The elapsed time (min:sec) since A23187 addition is indicated on each panel. The parasites shown here initiated egress around 2:09 for eGFP-AKMT-WT- and 9:19 for eGFP-AKMT-4A-expressing parasites. Fluorescence and DIC/fluorescence overlay images are shown. The red dotted circle in the overlay indicates untransfected Δ*akmt* parasites. (**c**) Bar-graph of percentage of vacuoles that egressed within ∼11 min of A23187 addition. The average is indicated next to the bar. The error-bars represent the standard deviation (SD). The number of individual dishes (n) and the total number of vacuoles (v) examined were indicated underneath the bars. The p-values from unpaired and one-tailed Student’s t-test were indicated above the bars; n.s., non-significance. (**d**) Box plot of the egress response time for vacuoles containing eGFP-AKMT-WT or eGFP-AKMT-4A transfected parasites that did egress. The central mark, upper and lower edges of the box, and upper and lower whiskers denote the median, 75th and 25th percentiles, and the maximum and minimum egress time respectively. The p-value from unpaired and one-tailed Student’s t-test was indicated above the boxes.

To test the effects of the dimerization-blocking mutations *in vivo*, we transiently expressed in Δ*akmt* parasites full-length AKMT-4A and AKMT-WT proteins with a fused eGFP tag at the N-terminus, to assess the localization and complementation efficacy of these alleles without long-term selection in a population (**Fig. 5b-d**). We found that, in intracellular parasites, the eGFP-AKMT-4A has a significantly more prominent cytoplasmic pool than eGFP-AKMT-WT (P <0.001, **Table 2 and Fig. 5b)**, suggesting that dimerization is important for efficiently targeting AKMT to the apical complex. Upon the addition of the calcium ionophore A23187, both eGFP-AKMT-WT and eGFP-AKMT-4A relocate from the apical complex (**Fig. 5b**). However, eGFP-AKMT-4A expressing parasites displayed a slower egress response to the A23187 stimulus than those expressing eGFP-AKMT-WT. As previously reported [6], the untransfected Δ*akmt* parasites had severe defects in A23187-induced egress (⟨%egress⟩ ≈ 4%). This defect was complemented by eGFP-AKMT-WT expression (⟨%egress⟩ ≈ 84%) and to a lesser extent by eGFP-AKMT-4A expression [⟨%egress⟩ ≈ 66%, P_(eGFP-AKMT-WT/ eGFP-AKMT-4A)_ ≈ 0.14, non-significance (n.s.)] (**Fig. 5c**). For those parasites that did egress, the ones expressing eGFP-AKMT-4A egressed with a marked delay compared with the ones expressing eGFP-AKMT-WT (**Fig. 5d**). The median length of time for the parasites to respond to the addition of A23187 to egress was ∼244 seconds for eGFP-AKMT-WT expressors, and ∼353 seconds for eGFP-AKMT-4A expressors, respectively (P ≈ 0.014). These results indicated that AKMT-4A is functionally compromised *in vivo*.

**Table 2.**
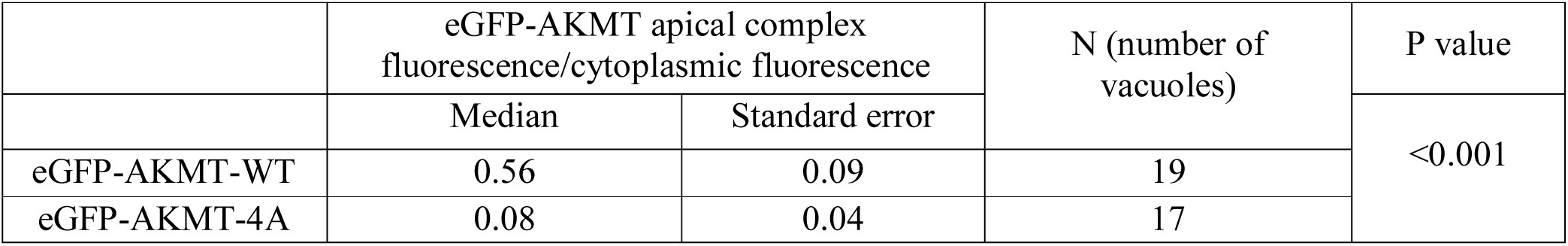
Relative distribution of eGFP-AKMT in the apical complex and cytoplasm in eGFP-AKMT-WT and eGFP-AKMT-4A expressing parasites.

|  | eGFP-AKMT apical complex<br>fluorescence/cytoplasmic fluorescence |  | N (number of<br>vacuoles) | P value |
| --- | --- | --- | --- | --- |
|  | Median | Standard error |  | <0.001 |
| eGFP-AKMT-WT | 0.56 | 0.09 | 19 |  |
| eGFP-AKMT-4A | 0.08 | 0.04 | 17 |  |

### AKMT is a unique homodimeric lysine methyltransferase

SET domains have been found in over a thousand proteins of diverse functions in organisms ranging from viruses to mammals [26]. Most SET domain-containing KMTs exist as monomers, but there are exceptions. Previous biochemical studies have demonstrated the self-association of two SET domains in human ALL-1 and in *Drosophila* TRITHORAX and ASH1 proteins [27]. Crystal structure of the viral SET (vSET) protein from the Paramecium bursaria chlorella virus, which has explicit methyltransferase activity for host histone H3 lysine 27, displayed a side-by-side homodimer [28] (**Fig. 6a**). The dimer was believed to be similar to that formed by the TRITHORAX and Ash1 proteins on the basis of the conserved interface residues among them [29]. However, in contrast to these dimeric KMTs, the dimerization of AKMT is distinct in several aspects. Firstly, the SET domain in AKMT is not involved in its dimeric interaction. Secondly, unlike the side-by-side arrangement in vSET, the dimerization of AKMT is via a C- and C-terminal (C-C) interaction mediated by the unique ASI motif together with the superhelical TPR-like CTD (**Fig. 6b**). Thirdly, the dimer of AKMT is structurally much more stable than that in the vSET structure, as indicated by the doubled BSA and the dramatically lower Δ^i^G in the AKMT dimer (**Fig. 6c**).

**Fig. 6.**
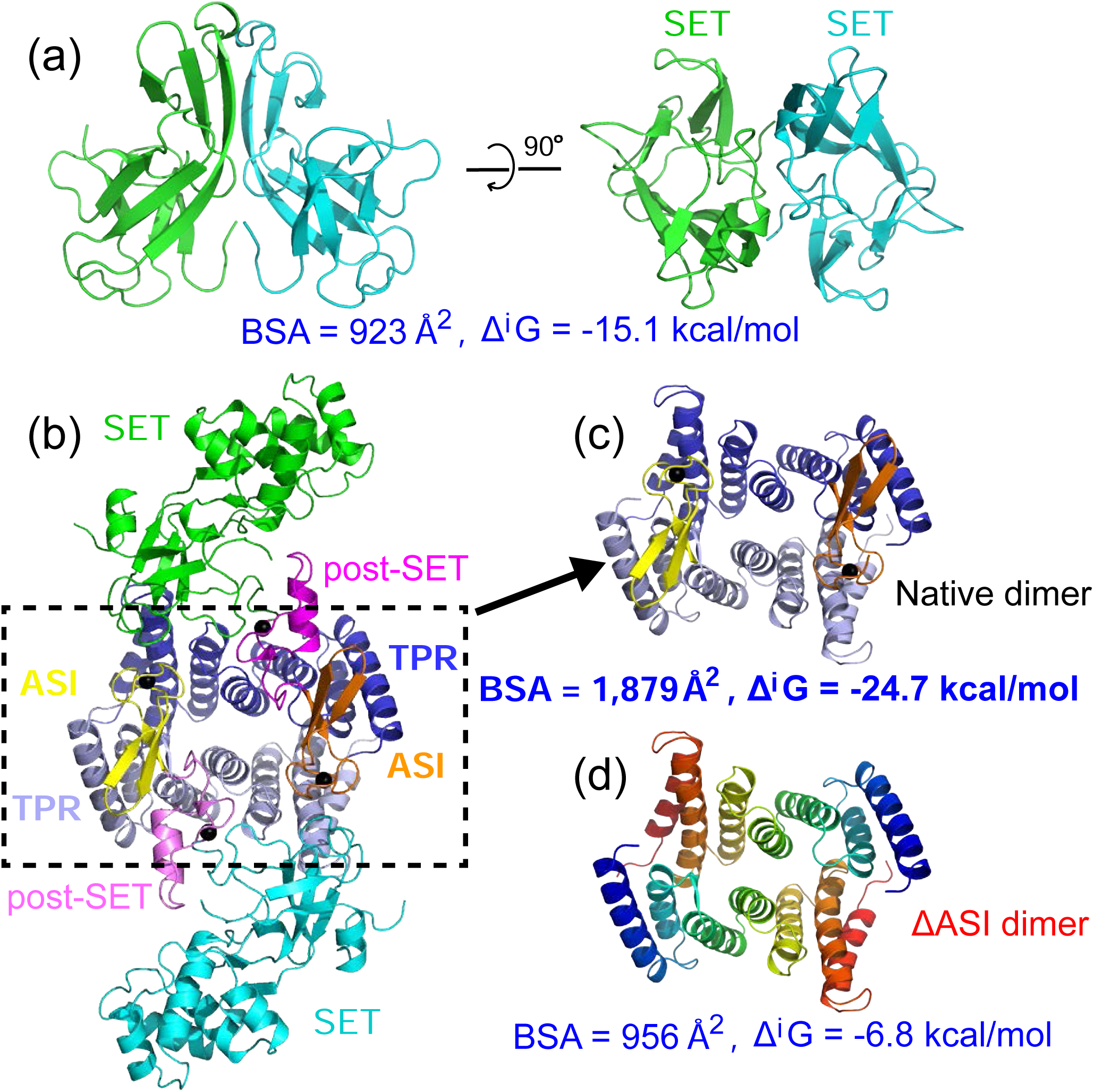
AKMT forms a novel homodimer mediated by both its ASI motif and TPR-like domain. (**a**) Ribbon diagram that shows the side-by-side arrangement of the SET domains in the vSET homodimer. (**b**) Crystal structure of the AKMT homodimer. (**c**) An extracted view of the dimeric interface formed by the ASI motif and the TPR-like domain of AKMT. (**d**) The same structure as in (c) but with both ASI motifs deleted. The two TPR-like domains are shown in rainbow coloration to demonstrate the extensive contacts across the two antiparallelly arranged structures. Notably, without ASI motifs, the buried interface area is reduced by ∼50%, and in the meantime Δ^i^G increases significantly, suggesting a substantial drop in structural stability.

Besides AKMT, there are many other unrelated proteins that also form homodimers via diverse interactions between two identical TPR or TPR-like domains. For examples, the anaphase-promoting complex/cyclosome (APC/C) subunit Cut9 from *Schizosaccharomyces pombe* forms an N- and N-terminal (N-N) intertwined dimer [30] (**Supplementary Fig. 7a**), the light chain of the mammalian microtubule motor kinesin-1 forms a C-C dimer [31] (**Supplementary Fig. 7b**), and the superhelical TPR domain of human O-linked GlcNAc transferase forms a middle-to-middle (M-M) dimer [32] (**Supplementary Fig. 7c**). However, the dimerization of AKMT is clearly different from all these reported TPR-mediated homodimers. Firstly, the TPR-like domain in the AKMT dimer is fully engaged to allow maximal interaction along the two conformationally complementary surfaces (**Fig. 6d**), unlike other TPR dimers mentioned above that are formed via the interaction of only parts of the two involved TPR domains (**Supplementary Fig. 7a-c**). Secondly, besides the TPR-like domain, the unique ASI motif is also heavily involved in the dimerization of AKMT. In fact, the two ASI motifs in the AKMT dimer not only provide a similar buried surface area to that of the TPR-like domains (**Fig. 6c and d**), but also contribute most of the hydrogen bonds and salt bridges between the two molecules in the complex (**Supplementary Fig. 4**). Deletion of the ASI motif would render the AKMT dimer significantly less stable (**Fig. 6c and d**). Therefore, AKMT is a unique homodimeric KMT that is stably maintained via an extensive network of interacting residues provided by both the ASI motif and the TPR-like domain.

### Structural comparison of AKMT with SMYD proteins

Previous phylogenetic analysis indicated that the AKMT and SMYD families branched off from a common node of SET domain-containing methyltransferases during evolution [22]. In the crystal structure, we found that AKMT shares an overall similar domain arrangement to SMYD proteins, including an N-terminal SET domain, a post-SET cysteine cluster, and a C-terminal TPR-like domain (**Fig. 7a**). In fact, many parts of AKMT, accounting for approximately 50% of the sequence in the crystal structure reported here, are very similar to the corresponding regions in SMYD proteins (**Supplementary Fig. 8**). Using the flexible alignment mode of the RAPIDO server [33], which is a tool for rapid alignment of protein structures in the presence of domain movements, the RMSDs of the locally aligned rigid bodies between AKMT and SMYD proteins were found to be approximately 1.08-1.34 Å (**Supplementary Fig. 8a**), which are nearly as good as those among different SMYD proteins that share much higher sequence identities with one another (30-35%) than with AKMT (∼18%). Superposition of the corresponding rigid bodies between AKMT and SMYD3 showed clear similarities of the structures (**Supplementary Fig. 8b-h**).

**Fig. 7.**
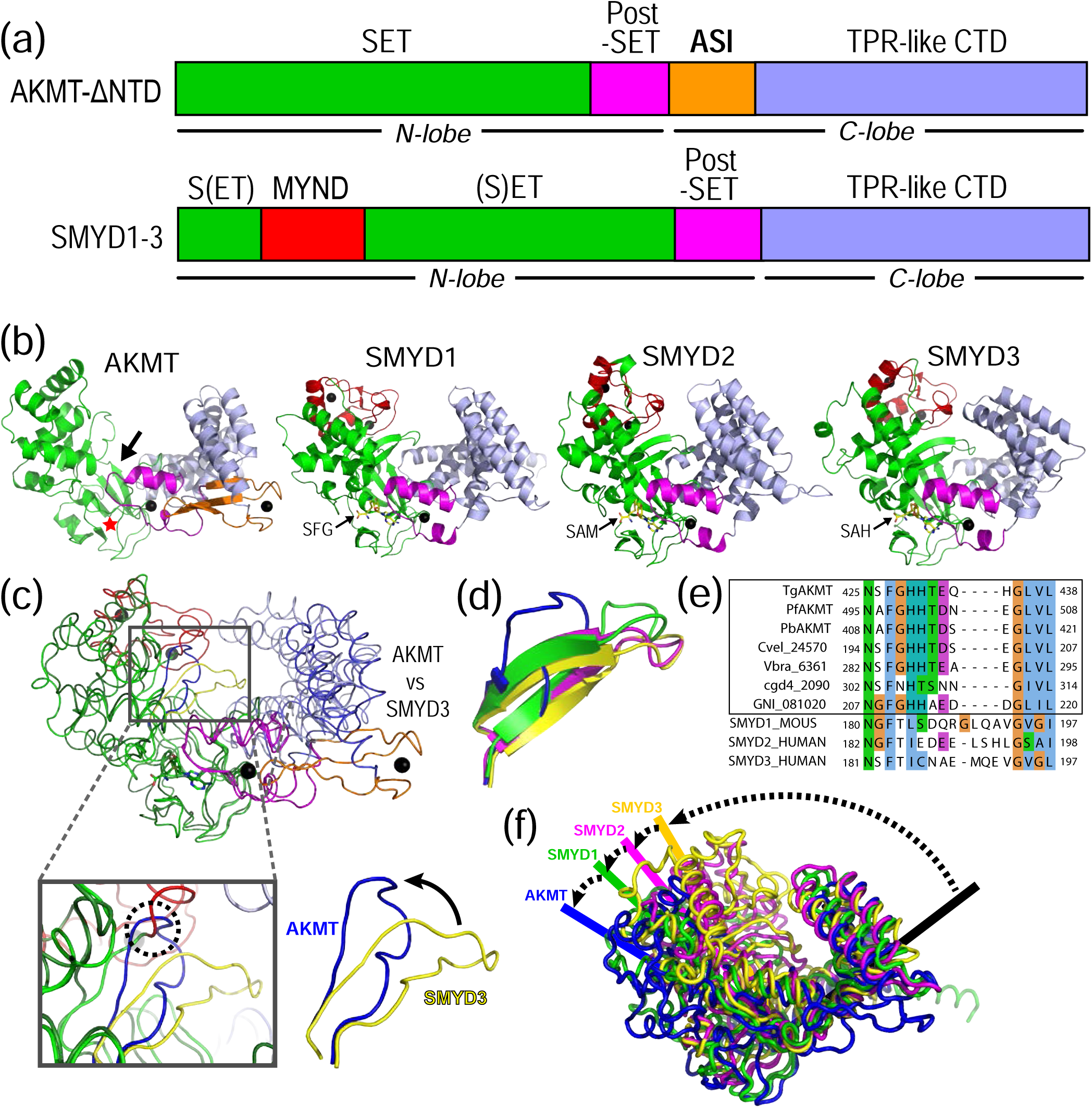
Structural comparison of AKMT with SMYD proteins. (**a**) Schematics of the domain organization of AKMT and SMYD1-3. (**b**) Ribbon diagrams of AKMT, SMYD1 (3N71.pdb), SMYD2 (5ARG.pdb), and SMYD3 (5EX0.pdb), in the same color scheme as in (A). All bound zinc ions are shown as black spheres. The putative cofactor and substrate binding sites in AKMT are marked by a red star and a black arrow, respectively. (**c**) Direct structural comparison between AKMT and SMYD3, with the “stirring” loop in AKMT colored in blue and the counterpart in SMYD3 in yellow. Superposition of AKMT on SMYD3 is based on the rigid body around their post-SET domains (see Fig. S7B-E). The “stirring” loops are better visualized in the enlarged view where potential clashes between the loop of AKMT and the MYND domain of SMYD3 are indicated by a dashed circle. The loops are also shown in isolation next to the inset to clearly illustrate the dramatic flip (black arrow). (**d**) Close-up view of the “stirring” loops in AKMT and SMYD1-3 structures, with regular β strands shown in flat sheets and irregular structures in loops. SMYD1, 2, 3, and AKMT are colored green, magenta, yellow, and blue, respectively. (**e**) Sequence alignment of the “stirring” loops of AKMT orthologs (boxed) and SMYD1-3. (**f**) Overlay of the AKMT and SMYD1-3 structures to demonstrate the variation of their inter-lobe grooves. The structures are superimposed on the C-terminal rigid body (see Fig. S7f-h).

Despite the high degree of local alignment with the SMYD proteins, AKMT has a number of unique structural features, which is consistent with a clear divergence between these two subfamilies after the speciation of the apicomplexan parasites [22]. First of all, the unique zinc-binding ASI motif in AKMT is not found in any SMYD proteins (**Fig. 7a, Supplementary Fig. 1b**). Secondly, AKMT and orthologs do not have the signature MYND (<u>My</u>eloid-<u>N</u>ervy-<u>D</u>EAF1) domain of SMYD proteins, which is a zinc finger motif embedded in the SET domain and located at the tip of the N-terminal lobe to gauge the entrance to the groove between the N- and C-terminal lobes (**Fig. 7b**). Previous studies suggested that the MYND domain serves as a protein-protein interaction module mainly in the context of transcriptional regulation [34, 35]. Lack of a MYND domain in AKMT might correlate with its specific function outside the nucleus [6].

Thirdly, the groove between the N- and C-terminal lobes in AKMT is considerably wider than that in all known SMYD structures, due to both the larger distance between its SET and TPR-like domains and the absence of MYND domain from its N-terminal lobe (**Fig. 7b**). The grooves in SMYD1-3 were proposed to accommodate their U-shaped substrates of various sizes [36]. Similarly, the larger open groove in AKMT may have evolved to target unique substrate(s) that regulates the motility switch of the parasite during egress. Such a variation in the open grooves of AKMT and SMYDs is coupled with a dramatic flip of a “stirring” loop (aa425-438 in AKMT) protruding out from the hinge connecting the N- and C-terminal lobes, close to the putative substrate binding site in AKMT (**Fig. 7c**). While the loops in all SMYDs adopt a similar conformation of a hairpin-like two-stranded antiparallel β sheet, the one in AKMT is irregularly folded and tilts towards the N-terminal SET domain (**Fig. 7d**). Sequence alignments show that the “stirring” loop in AKMT is not only 4-5 residues shorter than those in SMYD1-3 but also highly conserved in all AKMT orthologs (**Fig. 7e**), suggesting that the loop might play a unique and important role in AKMT function, such as to regulate the docking and/or selection of the so-far-unknown substrates. The difference of the putative substrate binding groove in AKMT and those in SMYD proteins is better viewed in the superimposed structures (**Fig. 7f**).

Finally, AKMT forms a distinct dimer that has not been seen in any SMYD proteins. This can be explained by the difference in the structural features at the C-terminal end of the proteins (**Fig. 8**). SMYD1-3 do not appear to possess the necessary features that are compatible for dimer formation. While the surface area at the C-terminal end of the AKMT structure is largely hydrophobic (**Fig. 8a**), the corresponding regions in the structures of SMYD1-3 are covered mostly by charged residues (**Fig. 8b-d**). Further, the buried interfaces on the two subunits of the AKMT dimer are also spatially and electrostatically complementary with each other, whereas those in SMYD1-3 structures do not support the docking of one molecule on top of the other to form a C- to C-terminal dimer as that of AKMT.

**Fig. 8.**
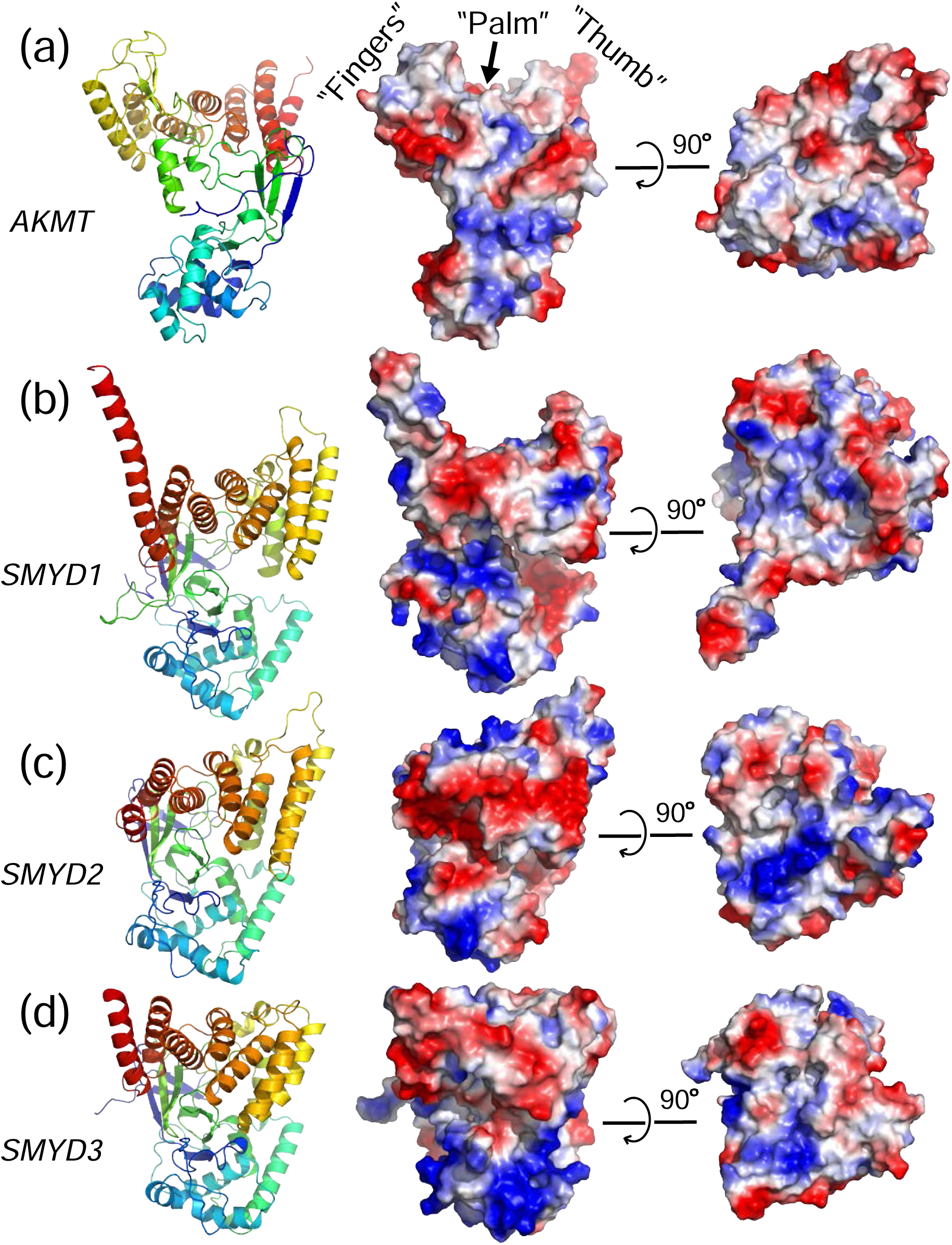
Comparison of dimeric AKMT with monomeric SMYD1-3. Two orthogonal views of the electrostatic plots of AKMT monomer (a), SMYD1 (b), SMYD2 (c), and SMYD3 (d). In contrast to the structurally and electrostatically complementary hand-like structure in AKMT, the corresponding surfaces in all SMYD proteins are unfavorable for intermolecular interactions. All plots were made using PyMOL (the PyMOL Molecular Graphics System, Version 2.0 Schrödinger, LLC).

## Discussion

In this work we describe our structural and biochemical characterizations of *T. gondii* AKMT. We discovered that the protein contains a number of structural features that are well-conserved among AKMT orthologs, but not found in other KMT families. In particular, the putative substrate binding groove between the N- and C-terminal lobes in AKMT is much wider than that in any known SMYD structures. Such a difference might reflect the need to accommodate different types of substrates. SMYD proteins regulate chromatin remodeling mainly by methylating histone lysines H3K4 (SMYD1-3) [37] and H3K36 (SMYD2) [38], both of which are located in the structurally disordered N-terminal tail of H3. It has been shown that SMYD proteins also methylate a number of non-histone targets, such as p53 [39], retinoblastoma tumor suppressor RB1 [40], MAP3K2 kinase [41], as well as vascular endothelial growth factor receptor-1 [42]. Similar to H3, targeted lysines in these non-histone proteins are positioned in either unstructured regions or extended loops. The disordered nature of targeted regions in these substrates allow them to accommodate a compact U-shaped structure to fit into the narrow substrate binding site between the N- and C-terminal lobes in the SMYD proteins [43-45].

In contrast, mass spectroscopy analysis of *Xenopus laevis* histone H3.3 methylated by AKMT *in vitro* revealed an unusual methylation site, K123, which is located at the C-terminal end of a 10-Å long α-helix right in the core of the folded structure [6]. This finding implies that in order to get access to residue K123 in H3.3, AKMT has to accommodate the α-helix containing it or even the whole folded structure into its substrate-binding site. It was recently reported that AKMT is required for recruiting a motility-relevant protein (the glideosome-associated connector, GAC) to the apical localization, and depletion of AKMT led to the disappearance of several large proteins in a western blot using α-H4K20Me3 antibodies to detect methylated lysine residues [46]. The identities of these high molecular weight proteins and whether they are true AKMT substrates are still yet to be determined. Nevertheless, it is conceivable that the wide-open groove leading to the putative substrate binding site in AKMT might be necessary for AKMT to accommodate either a large substrate or its subdomains (up to 30 Å in diameter), or several substrates with various sizes and shapes, in which case a wider groove would provide more flexibility for AKMT to target different substrates.

In order to stabilize such an open conformation, AKMT seems to have evolved to adopt a homodimeric conformation. Disrupting the dimerization reduces the thermostability of the protein, and dropped the Tm by ∼8°C (**Fig. 4j**). AKMT dimerization might also cooperatively regulate cofactor and/or substrate binding to affect enzymatic activity. Indeed, our ITC data demonstrate that, while the wild-type and the mutant AKMT bind cofactors with affinities at the same order of magnitude, the monomeric mutant (AKMT-ΔNTD-4A) did show consistently lower binding affinity for SFG than the dimeric truncation (AKMT-ΔNTD) (**Fig 4b**).

Consistent with the biophysical measurements, our *in vitro* enzymatic activity assays, as well as *in vivo* egress rescue tests, revealed that the mutant with an impaired dimerization interface (*i.e.* mutant AKMT-4A) is less active than the wild-type and results in delayed egress (**Fig. 5**). It is worth noting that while a delay in egress does not significantly impact the parasite survival under the lenient culture conditions in the laboratory, in an animal host, this delay may render the parasite vulnerable to the attack from the immune system and can be lethal to the parasite. For instance, the loss of perforin (a pore-forming protein secreted by the parasite during egress to disrupt the membrane of the parasitophorous vacuole) delays parasite egress, but does not have any apparent impact on parasite growth in tissue culture [47]. However, it completely abolishes parasite virulence in mouse infections.

While the AKMT-4A mutant displays defects in *vivo* and *in vitro,* it is not a null mutant. This is possibly because the R521A/R672A/D676A/N679A mutations only partially disrupt the dimerization of AKMT, as the AKMT-ΔNTD dimer is maintained by a large number of other interactions besides the hydrogen bonds and salt bridges disrupted in the monomeric mutant (**Fig. 3e)**. Indeed, size exclusion chromatography (SEC) analyses of AKMT-ΔNTD-4A showed that a small fraction of the protein eluted as dimers at >10 mg/ml (data not shown). The *ab initio* shape reconstructed with SAXS data for AKMT-ΔNTD-4A at a similar concentration in a batch measurement displayed an elongated “tail” connected to the spherical envelope that is comparable to the AKMT monomer (**Supplementary Fig. 9a**), while the SEC-SAXS data did not show such extra “tail” in the *ab initio* model (**Supplementary Fig. 9b**). It suggests that while the majority of AKMT-ΔNTD-4A protein forms monomers, a small fraction could form dimers, or else there is a quick exchange between monomer and dimer at an elevated concentration. Furthermore, as mentioned above, mutations of any of the four conserved cysteine residues in the ASI motif resulted in insoluble proteins, possibly due to the deleterious effect caused by the complete disruption of the dimeric interaction. These observations suggest that the 4A mutant probably only partially disrupts the dimeric conformation of AKMT. However, we also cannot exclude the possibility that the monomeric form of AKMT is partially active.

Besides the folded region of AKMT that has been extensively discussed above, AKMT additionally contains a structurally disordered N-terminal region (aa1-290) (**Supplementary Fig. 1a and b**). Such an unstructured segment does not exist in any of the closely related SMYD proteins from animals. Previous studies on different AKMT truncations demonstrated that the N-terminal region makes only minor contributions to the specific cellular localization or the enzymatic activity of the protein [22]. Nevertheless, given that there are so many negative charges spreading throughout the N-terminal sequence (31% of aa1-290 in TgAKMT are D or E) (**Supplementary Fig. 1b**), the unstructured N-terminal region of AKMT may play subtle regulatory roles in fine-tuning the function of the enzyme, which needs to be addressed in future studies.

The protein sequences for the distinct structural features of TgAKMT are highly conserved among the apicomplexans and their free-living relatives, but not found in animals. Future identification of AKMT substrates and structural studies of AKMT-substrate complex will shed light on the specificity of substrate binding for designing selective drugs and for informing the evolution of the function of the SMYD related KMTs. High-resolution structures of AKMT in complex with cofactors should also provide structural information about the shape and chemical environment of the cofactor-binding site, which could be potentially used for the development of the cofactor-competitive selective inhibitors.

## Materials and methods

### Cloning and site-directed mutagenesis

Sequence encoding full-length *T. gondii* AKMT (TGGT1_216080) was amplified from cDNA using PCR and ligated into the pET15b vector (Novagen) between NdeI and BamHI sites at the 5’ and 3’ ends, respectively. The plasmid when expressed provides an N-terminal His_6_ tag followed by a thrombin cleavage site prior to the target protein. All truncations of AKMT were subsequently generated in a similar manner using the precloned full-length AKMT construct as the PCR template. The AKMT-ΔNTD fragment was amplified using 5’-gcagctgcatatggccaaccccttagctccatac-3’ (forward) and 5’-gaccggatcctcaactggccggtaggcg-3’ (reverse) primers and ligated into pET15b to produce pET15b-His6-AKMT-ΔNTD (aa289-709). The AKMT-ΔNTD-short insert was generated using the same reverse primer and the forward primer 5’-gcagctgcatatggtgccaggcaaggggcgatg-3’ and ligated into pET15b to produce pET15b-His6-AKMT-ΔNTD-short (aa309-709). The AKMT-ΔNTD-4A mutant was generated in two steps. First, three mutations R672A/D676A/N679A were introduced by Megaprimer PCRs [48]. Forward primer 5’-ctgaacgtgatatctgcacactttgaggccgcgtatgctcttctgtac-3’ and reverse primer 5’-gaccggatcctcaactggccggtaggcg-3’ were used for generating the 142-bp megaprimer that was subsequently elongated in a second PCR with the same primers used for AKMT-ΔNTD amplification. In the second step, the fourth mutation, R521A, was introduced using a site-directed mutagenesis protocol (forward primer: 5’-caacgcccgagggtttgcatgccctctgtgtg-3’, reverse primer: 5’-cacacagagggcatgcaaaccctcgggcgttg-3’) to generate pET15b-His6-AKMT-ΔNTD-4A (aa289-709, R512A/R672A/D676A/N679A). All mutations were verified by DNA sequencing.

The plasmid pmin-eGFP-AKMT-4A was constructed using NEBuilder HiFi DNA assembly kit (NEB cat# E5520S) with the following three components: (1) the pmin-eGFP vector backbone released from pmin-eGFP-AKMT (aa1-300) [22] by BglII-AflII digestion; (2) PCR product that contained AKMT (aa1-298) sequence and appropriate overlapping regions for Gibson assembly [amplified using pmin-eGFP-AKMT as the template and 5’-ctgtacaagtccggactcagatct-3’ and 5’-gcagtgtgtatggagctaaggggttggc-3’as the primers]; (3) PCR product that contained AKMT (aa299-709)-4A sequence and appropriate overlapping regions for Gibson assembly [amplified using pET15b-His_6_-AKMT-ΔNTD-4A as the template and 5’-gggcagcttctgtttacttaagtcaactggccggtaggcgttcctcg-3’ and 5’-ttagctccatacacactgccccagatt-3’ as the primers].

### Protein expression and purification

For purifying recombinant proteins, pET15b-His_6_-AKMT (aa1-709), pET15b-His_6_-AKMT-ΔNTD (aa289-709), pET15b-His_6_-AKMT-ΔNTD-4A (aa289-709, R512A/R672A/D676A/N679A) and pET15b-His_6_-AKMT-ΔNTD-short (aa309-709) constructs were used to transform competent *E. coli* BL21(DE3) cells. Bacterial cells were grown in LB medium containing 50 μg/ml ampicillin at 37 °C until their OD_600_ reached 0.6-1.0 (approximately 2-3 h). Protein expression was induced using 250 μM of isopropyl-beta-D-thiogalactopyranoside (IPTG), and the cells were then incubated at 18°C overnight. Cells were harvested by centrifugation (6,000×g, 12 min, 4°C). Cell pellets were resuspended in prechilled lysis buffer [20 mM Tris-HCl pH 8.0, 100 mM NaCl, 20 mM imidazole, 10 mM beta-mercaptoethanol, 5% (v/v) glycerol] and were lysed using an EmulsiFlex-C3 homogenizer (Avestin). Cell debris was removed by centrifugation (25,000×g, 40 min, 4°C), and the supernatant was filtered through a 0.45-μm pore size filter and then loaded onto a 5-ml Ni-HiTrap column (GE Healthcare) pre-equilibrated in lysis buffer. After washing with 5 column volumes (cv) of lysis buffer, bound protein was eluted using a linear gradient concentration of imidazole in the lysis buffer (20 to 600 mM, 10×cv). The N-terminal His_6_ tag was removed by incubating the purified protein with ∼5% (w/w) thrombin (4°C, overnight). In order to achieve higher purity, all proteins were subjected to SEC using a Superdex S-200 16/60 column (GE Healthcare) pre-equilibrated with running buffer (20 mM Tris-HCl pH 8.0, 100 mM NaCl). Purified proteins were concentrated using Amicon Ultra Centrifugal Filter Units (Millipore) with appropriate molecular weight cutoffs.

For purifying proteins used in the protein lysine methyltransferase (PKMT) assay, BL21-CodonPlus(DE3)RP strain *E. coli* (CAT#230255: Stratagene, CA) were transformed using pET15b-His6-AKMT-ΔNTD or pET15b-His6-AKMT-ΔNTD-4A, and grown and induced as described above. 50 ml of bacterial culture was pelleted (6,000×g, 12 min, 4°C) and stored at -80 °C until use. Bacterial pellets were resuspended and lysed in Tris-acetate lysis buffer (8mM Tris-acetate pH 7.5, 7 mM Tris base pH unadjusted, 100 mM KAcetate, 1mM MgAcetate) containing 1 μM TAME (CAT# T4626: Sigma), 1 μM PMSF (CAT# P7626: Sigma), 1 μM Pepstatin A (CAT#P5318, Sigma), 0.5% Triton-100 (TX-100) and 25 mg/ml CelLytic express (CAT# C1990: Sigma), followed by sonications (Branson Sonifier 250). His_6_-AKMT bound to Talon-resin (CAT# 635503: Clontech) for 1 hr at 4°C was washed with Tris-acetate lysis buffer containing 5 mM imidazole-acetate, and eluted with Tris-acetate lysis buffer containing 200 mM imidazole. dithiothreitol was added to the eluted proteins to 4 mM. Eluted proteins were dialyzed against dialysis buffer (Tris-Acetate lysis buffer containing 4 mM dithiothreitol) at 4°C in Slide A lyzer Mini Dialysis Devices (CAT# 88401: Thermo Scientific) with 3 buffer exchanges (3 to 15 hr between each exchange). For long-term storage at -20°C, glycerol was added to dialyzed proteins to a final concentration of 36% (v/v). Protein concentrations were measured using a dual-absorbance Bradford assay [49].

### Crystallization and data collection

Initial trials to crystallize all the three AKMT truncations (aa289-709, aa296-709, and aa301-709) did not yield any crystals. In order to facilitate the crystallization process we carried out reductive lysine methylation on the purified proteins using a published protocol [50]. One of the truncations (aa289-709, denoted as AKMT-ΔNTD) with methylated surface lysine residues was successfully crystallized using the hanging drop vapor diffusion method against a reservoir solution containing 0.1 M Tris-HCl pH 9.0, 0.2 M MgCl_2_, 30% (w/v) polyethylene glycol (PEG) 4,000 (or 20 to 30% PEG 8,000). For harvesting, crystals were soaked in the same reservoir solution augmented with increasing concentrations of glycerol (final concentration 20% [v/v]), mounted in loops (Hampton research), and flash frozen in liquid nitrogen. X-ray diffraction data collection was carried out at the European Synchrotron Radiation Facility (ESRF) on the beamline ID29. A complete and highly redundant data set to 2.1-Å resolution was collected at the absorption edge of zinc (λ = 0.9792 Å). The crystals belonged to the space group P1.

### Structure determination and analyses

Diffraction data were integrated and scaled using XDS [51]. Phases were determined *de novo* using the single-wavelength anomalous dispersion (SAD) technique based on the anomalous signal from the zinc atoms bound to AKMT. In total, eight zinc ions were located and initial experimental maps were calculated using AutoSol in the software suite Phenix [52]. A preliminary structural model was built using AutoBuild, and the resulting model was optimized by multiple rounds of manual rebuilding in COOT [53] and refinement in Phenix [52]. The final model was validated in MolProbity [54].

Detailed analysis of intermolecular interfaces was performed using PISA software [23]. Structural alignment of AKMT with SMYD1-3 proteins was done by the RAPIDO server [33].

### Static light scattering (SLS)

SLS measurements were carried out by coupling SEC with mass determination. 50 μl of protein sample at 2 mg/ml was analyzed on a Superdex S-200 10/300 GL column (GE Healthcare) pre-equilibrated with a buffer containing 20 mM Tris-HCl pH 8.0, 100 mM NaCl, 1mM dithiothreitol, and 1% (v/v) glycerol. The sample was run at a flow rate of 0.5 ml/min on a High performance liquid chromatography (HPLC) system (Agilent Technologies 1260 infinity) which was directly connected to a Mini-DAWN Treos light-scattering instrument (Wyatt Technology Corp., Santa Barbara, CA). Data analyses were carried out using ASTRA software provided by the manufacturer. Final curves were built in SigmaPlot [55].

### Circular dichroism (CD)

Far-UV CD spectra of AKMT-ΔNTD and AKMT-ΔNTD-4A between 180 and 280 nm were measured on a Chirascan plus CD spectrometer (Applied Photophysics) in a cuvette with a 0.5-mm path length. Proteins were diluted to approximately 0.2 mg/ml using a buffer containing 10 mM Tris-HCl (pH 8.0) and 100 mM NaF. Data points were corrected for buffer signal and drifts. CD curves were generated using SigmaPlot by averaging data collected from two scans for each protein sample.

### Differential scanning fluorimetry (DSF)

DSF measurements were performed on a CFX Connect Real-Time PCR machine (Bio-rad). Proteins were diluted to 0.4 mg/ml using a buffer containing 10 mM Tris-HCl pH 8.0 and 100 mM NaCl. To prepare for the measurements, 24 μl of the protein sample was mixed with 1 μl of 20 × SYBR orange dye (ThermoFisher Scientific). Each sample was prepared and measured in duplicates, and the two measurements were averaged to produce the final curve. Melting temperature (T_m_) values were obtained using the software provided by the manufacturer.

### Isothermal titration calorimetry (ITC)

ITC measurements were performed on a MicroCal™ iTC200 microcalorimeter (GE Healthcare). Cofactors SAM and SFG (Sigma-Aldrich) were dissolved in a buffer containing 20 mM Tris-HCl (pH 8.0) and 100 mM NaCl. Protein samples were dialyzed overnight against the same buffer before the measurements. A typical ITC titration experiment consisted of 20 injections of the cofactor solution, first with 1 × 0.2 μl and then with 19 × 2 μl, into the cell filled with 200 μl of protein solution. All measurements were carried out under constant stirring at 350 rpm, and each injection lasted for 4 sec with a 180 sec interval between two injections. Titration peaks were analyzed using the Origin 7.0 Microcal software and corrected for the SAM/SFG dilution heat measured by injecting cofactors into the buffer containing no protein using the same protocol described above. Non-linear least-squares fitting using one binding site model was used to calculate the association constant (K_a_) and stoichiometry values. Dissociation constants (K_d_) were calculated according to the formula K_d_ = 1/K_a_.

### Denaturing liquid chromatography-mass spectrometry (LC-MS)

The purified sample of AKMT-ΔNTD at 1mg/ml was reduced by incubating with 100 mM dithiothreitol (30 min, RT). HPLC was performed on a Dionex Ultimate 3000 HPLC system configured with the Chromeleon 6.0 software (both Thermo Fisher Scientific). The protein sample was applied on an Aeris Widepore C4 column and run at a flow rate of 300 μl/min using a 6-min step gradient with increasing acetonitrile (ACN) concentration from 9 to 36% (w/v) in 5 min, and then from 36 to 63% (w/v) in 1 min at 50°C. The HPLC system was coupled online to a quadrupole-time of flight mass spectrometer Synapt G2-Si via a Z Spray ESI source (both Waters) operated via the MassLynx V 4.1 software package (Waters). Mass spectra were acquired in the m/z range from 500-2000 Th at a scan rate of 1 sec. Glu[1]-Fibrinopeptide B (Glu-Fib) was used as a lock mass and spectra were corrected during data acquisition. Acquired data were analyzed with the MaxEnt 1 algorithm to reconstruct the uncharged average protein mass.

### Native mass spectrometry

The protein sample at 22 mg/ml was incubated with 2 mM SFG (1 h, 4°C). After incubation, the sample was diluted to a final protein concentration of 2 mg/ml. The original buffer (20 mM Tris-HCl, pH 8.0 and 100 mM NaCl) was substituted for 400 mM ammonium acetate at pH 8.0 in Bio-Spin 6 columns (Bio-Rad Laboratories). The protein solution was transferred into pre-opened metal-coated emitters (PicoTips™, New Objective) and measured off-line on the Synapt G2-Si using a source voltage of 1.9 kV. Mass spectra were acquired in the m/z range from 100-5000 Th at a scan rate of 1 sec. To facilitate efficient desolvation of the complex, the trap collision energy was set to 12 eV and was increased up to 100 eV to shake off the non-covalently bound SFG molecules. Mass determination was performed manually.

### Small angle X-ray scattering (SAXS)

Scattering curves for AKMT-ΔNTD were collected at the beamline BM29 Bio-SAXS at the ESRF in Grenoble, France, using an online SEC setup. The protein sample (200 μl, 5 mg/ml) was applied to a Superdex-200 10/30 column (GE Healthcare) equilibrated with 20 mM Tris-HCl (pH 7.5), 150 mM NaCl, 1 mM TCEP [tris(2-carboxyethyl)phosphine] and 1% (v/v) glycerol. SAXS data frames were recorded every second during the chromatography run. The data were processed in PRIMUS [56]. Fifty buffer frames selected after the protein elution peak were averaged and used for buffer subtraction from 100 averaged protein frames from the peak region where R_g_ value was determined to be stable. The program GNOM [57] was used to plot the pair distribution function and to determine the D_max_ value of the protein from the scattering profile. An *ab initio* model of the AKMTΔNTD dimer was reconstructed by running 10 cycles of dummy beads model building and simulated annealing implemented in DAMMIF [58]. The most probable models were selected and averaged in DAMAVER [59]. Superposition of the *ab initio* model with the crystal structure was performed by SUPCOMB [60]. SAXS curves for A-B and A-C dimers of the AKMT-ΔNTD were calculated and fitted to the experimental data using CRYSOL [61].

### Protein lysine methyltransferase (PKMT) assay

The PKMT assay was performed as described previously [6] with the following modifications. For activity assays with recombinant wild-type and mutant AKMT truncations (AKMT-ΔNTD and AKMT-ΔNTD-4A, respectively), reactions were set up with 120 nM to 15 nM of AKMT truncations, 4.1 μM human histone H3.3 (CAT# M2507S: New England Biolabs), 0.75 mM dithiothreitol, 1.66 μCi (9.2 μM) ^3^H-S-adenosyl-L-methionine (^3^H-SAM) (CAT# NET155001MC: Perkin Elmer) in Tris-acetate lysis buffer (see above). 0.1mg/ml BSA (NEB B9001S) was included in each reaction to prevent non-specific sticking. Reactions were set up on ice and incubated in a room temperature water bath for 0, 2.5, 4.5, 8.5 and 16.5 min. At each timepoint, aliquotes of eight 12 μl reaction mix containing 120, 60, 30, 15 nM AKMT-ΔNTD-4A or AKMT-ΔNTD were transferred by an eight channel pipetman to a 0.45μm nitrocellulose membrane (CAT#10600002: GE Healthcare Life Sciences) assembled into a 96 well microsample filtration manifold (CAT#SRC096/0: Schleicher & Schuell), absorbed to the membrane by vacuum assisted filtration, and washed several times by ∼180 μl DPBS. The membrane was then air-dried and exposed to a tritium phosphor screen (CAT#28956482: GE Healthcare Life Sciences) for ∼ 22 hr, which was then scanned by Typhoon FLA 9500 (GE Healthcare Life Sciences) with 50 μm resolution. To visualize total proteins, the membranes were first rehydrated in DPBS, stained with amido black staining solution [0.1% (w/v) amido black 10B, 45% (v/v) methanol, 10% (v/v) acetic acid] for ∼5 min and rinsed briefly 4-6 times in the destaining solution [5% (v/v) methanol and 7% (v/v) acetic acid]. For quantification, the signal from the Typhoon scan of the phosphorimager was first normalized against the corresponding protein staining by amido black to correct for variations in protein retention on the dot blot (“retention corrected tritium signal”). Individual retention corrected tritium signals were then normalized against the sum of corrected signals of reactions in the same experiment to generate “normalized tritium signal” (Y-axis, **Fig. 5a** graph). Three independent experiments were conducted. The error bars in **Fig. 5a** graph represent ±2 standard deviations.

### *Toxoplasma* and host cell cultures, and parasite transfection

Tachyzoite *T. gondii* Δ*akmt* parasites (RhΔhx strain) were maintained by serial passage in confluent human foreskin fibroblast (HFF) (CAT# SCRC-1041: ATCC) monolayers in Dulbecco’s Modified Eagle’s Medium (CAT# 10569-010: Life Technologies-Gibco), supplemented with 1% (v/v) heat-inactivated cosmic calf serum (Cat# SH30087.3: Hyclone) as previously described [62, 63]. *T. gondii* transfections were carried out as previously described [64].

### Egress assay

Plasmids of pmin-eGFP-AKMT-4A or pmin-eGFP-AKMT-WT [6] were used to transfect Δ*akmt* parasites as previously described [6]. The parasites were added to 35 mm dishes (#1.5) with a 20 mm microwell (CAT# P35G-1.5-20-C: MatTek, Ashland, MA) seeded with a near confluent monolayer of HFF or A7r5 (CAT# CRL-1444: ATCC) grown for ∼48 h. Before imaging, the culture medium was replaced with 1 ml of phenol red-free CO_2_-independent medium (SKU#RR060058: Life Technologies-Gibco) supplemented with 1% (v/v) heat-inactivated cosmic calf serum, GlutaMAX^®^(SKU#35050-061: Life Technologies-Gibco), and sodium pyruvate (SKU#11360: Life Technologies-Gibco). 0.5 ml of 15 μM A23187 in CO_2_-independent medium was added (for a final A23187 concentration of 5 μM) to induce egress. Egress induction and imaging collection were carried out at 37°C on a DeltaVision imaging station (GE Healthcare-Applied Precision, WA) constructed on an Olympus IX-70 inverted microscope base. The exposure time was 0.2 sec in the GFP channel for all images. Differential interference contrast (DIC) and GFP fluorescence images were recorded. Vacuoles from 6-7 independent transfections in 12, 15 and 13 individual dishes were performed on eGFP-AKMT-WT and eGFP-AKMT-4A expressing and parental Δ*akmt* parasites respectively. The p-values in **Fig. 5c** and **Fig. 5d** were calculated by unpaired and one-tailed Student’s t-test.

### Quantification of AKMT distribution

To quantify the difference in the distribution of eGFP-AKMT-WT and eGFP-AKMT-4A, image stacks of vacuoles with 8 or fewer parasites collected prior to A23187 treatment in the above egress assays were used (17 vacuoles for eGFP-AKMT-4A expressing parasites and 19 vacuoles for eGFP-AKMT-WT expressing parasites). Images were analyzed in a double-blind test. Pre-processing consisted of first correcting image intensities for non-uniform illumination by ‘flat-fielding’ (i.e. dividing each image by the image of a uniformly fluorescent specimen that had been normalized to 1.0 at its peak), and in addition the region for analysis was restricted to the portion of the image where illumination was at least 85% of its maximum. Next, the nominal exposure time for each image was adjusted to compensate for differences in actual illumination intensity during each individual image acquisition, as recorded by photosensor hardware and software built in to the DeltaVision system. Each image was then divided by its adjusted exposure time to yield normalized intensities in units of grey-levels/sec. Single optical sections at optimal focus were used for analysis, drawn from 3D stacks of 6-10 z-slices.

Each apical complex was marked with a cursor on a screen display of the image, after which its exact x,y,z, location was found by searching within a small x-y neighborhood of the cursor mark, through all z-planes. Intensity within a circular ROI with diameter twice the size of the PSF (N.A 1.4 objective lens, 520 nm) was summed and corrected for local background.

Total fluorescence within the parasite was measured by summing the intensities within a manually specified elliptical ROI that enclosed the entire parasite, including the apical complex, and then correcting this sum for local background. From this total, the previously measured apical complex value was subtracted to yield the fluorescence in the cytoplasm only. If daughters were present, their apical complexes were measured as above, and their summed intensity was also subtracted from the total parasite fluorescence.

The ratio of the apical complex and cytoplasmic fluorescence is then computed as a measure of relative distribution of AKMT in eGFP-AKMT-4A and eGFP-AKMT-WT expressing parasites. Data points within 4 standard deviations of the median were used to compute the p-value by unpaired and one-tailed Student’s t-test (**Table 2**).

All image analysis was carried out with an updated version of the Semper software package (source code generously provided by Dr. Owen Saxton) [65].

## Accession code

Coordinates and structure factors have been deposited in the Protein Data Bank (PDB) under accession code 6FND.

## Acknowledgements

The authors are grateful to the staff at the beamlines of ID29 and BM29 at the European Synchrotron Radiation Facility (ESRF) for their help with X-ray diffraction and SAXS data collections. We also thank the staff at the mass spectrometry facility of the MFPL for their assistance in our mass spectrometric measurements. We greatly appreciate Dr. Brooke Morriswood for critical reading of the manuscript and Dr. John M. Murray for carrying out a double-blind image analysis. This work was supported by the grant P23440-B20 from the Austrian Science Fund (FWF) to GD, and funding from the National Institutes of Health/National Institute of Allergy and Infectious Diseases (R01-AI098686) and the March of Dimes (6-FY15-198) awarded to KH. YP is enrolled in the Integrative Structural Biology PhD program funded by the FWF.

## Author Contributions

G. Dong and K. Hu conceived the project. Y. Pivovarova and G. Dong designed the experiments and performed analyses for all structural studies. J. Liu and K. Hu conducted the *in vitro* methylation and *in vivo* egress assays. J. Lesigang, O. Koldyda, and R. Rauschmeier assisted in generating preliminary data to prove the feasibility of the project. Y. Pivovarova, J. Liu, K. Hu, and G. Dong wrote the manuscript.

## Abbreviations

AKMT: <u>a</u>pical complex lysine (<u>K</u>) <u>m</u>ethyltransferase
ASI: <u>A</u>KMT-<u>s</u>pecific insertion
KMT: lysine methyltransferase
MYND: <u>my</u>eloid-<u>N</u>ervy-<u>D</u>EAF1
SET: <u>s</u>uppressor of variegation [Su(var)3-9], <u>E</u>nhancer of zeste [E(z)] and <u>T</u>rithorax
TPR: tetratricopeptide repeat
SMYD: SET-and myeloid-Nervy-DEAF-1 (MYND)-domain

